# Post-translational digital data encoding into the genomes of mammalian cell populations

**DOI:** 10.1101/2024.05.12.591851

**Authors:** Alec Callisto, Jonathan Strutz, Kathleen Leeper, Reza Kalhor, George Church, Keith E.J. Tyo, Namita Bhan

## Abstract

High resolution cellular signal encoding is critical for better understanding of complex biological phenomena. DNA-based biosignal encoders alter genomic or plasmid DNA in a signal dependent manner. Current approaches involve the signal of interest affecting a DNA edit by interacting with a signal specific promoter which then results in expression of the effector molecule (DNA altering enzyme). Here, we present the proof of concept of a biosignal encoding system where the enzyme terminal deoxynucleotidyl transferase (TdT) acts as the effector molecule upon directly interacting with the signal of interest. A template independent DNA polymerase (DNAp), TdT incorporates nucleotides at the 3’ OH ends of DNA substrate in a signal dependent manner. By employing CRISPR-Cas9 to create double stranded breaks in genomic DNA, we make 3’OH ends available to act as substrate for TdT. We show that this system can successfully resolve and encode different concentrations of various biosignals into the genomic DNA of HEK-293T cells. Finally, we develop a simple encoding scheme associated with the tested biosignals and encode the message “HELLO WORLD” into the genomic DNA of HEK-293T cells at a population level with 91% accuracy. This work demonstrates a simple and engineerable system that can reliably store local biosignal information into the genomes of mammalian cell populations.

## Introduction

As the medium of information exchange and heredity in biological systems, DNA evolved to be an elegant information storage substrate, capable of information densities exceeding 200 Pbytes/g^1^. While organisms encode peptide sequences and regulatory topologies, among other data, in their genomes, biological systems can also be repurposed to write data into cellular DNA for technological applications. A variety of enzymatic data encoding systems have been applied to DNA-based data storage and cellular biosignals. Generally, these systems employ recombinases^2,3^, nucleases^4–7^, base editors^8^, or spacer acquisition enzymes^9–12^ as the effector molecules encoding the signal of interest to predetermine loci on plasmid or genomic DNA. Often, these systems encode information with defined changes to DNA sequences in response to inputs of interest^13^, however some systems instead use the accumulation of stochastic mutations^5,6,14^ or insertions^9^ at said loci. Recently, we demonstrated an alternative biorecording modality *in vitro* that encodes information in the overall distribution of bases in DNA rather than as defined sequences^15^. In this modality, data is not encoded in the precise sequence of bases, but by the average composition in a stretch of bases. For example, an AT-rich stretch of DNA might encode a different signal state than a GC-rich stretch of DNA. To achieve this, we used a template-independent DNA polymerase, terminal deoxynucleotidyl transferase (TdT) as the effector molecule. During *in vitro* single-stranded DNA (ssDNA) synthesis, relevant signals alter nucleotide selectivity of TdT, thus encoding sequential signal changes in the nucleotide composition along the synthesized ssDNA.

Here, we demonstrate the feasibility of adapting this DNA composition-based system for encoding information in living mammalian cell populations. By employing self-targeting CRISPR-Cas9 (also known as homing guide RNA, hgRNA) system^6,14^ to generate multiple and continuous double-stranded breaks at randomly distributed genomic hgRNA sites, we are able to provide the 3’OH ends needed as substrate for TdT. TdT-catalyzed untemplated ssDNA is synthesized at these hgRNA sites during each round of Cas9 induced DNA break and repair cycle. Much like *in vitro* extensions with TdT, we observed that the DNA synthesized at such genomic sites exhibited a distinct, reproducible composition of bases. Further, the perturbation of the intracellular nucleotide pools by treatment with different nucleosides resulted in altered distribution of bases incorporated by TdT at these hgRNA sites. Six different cellular treatment conditions that resulted in statistically distinguishable base distributions at hgRNA sites were identified.

To demonstrate that these differences in DNA composition can be used to encode digital data, we developed a simple alphanumeric encoding scheme. We treated pairs of barcoded hgRNA expressing cell populations with nucleosides at various concentrations to encode data. The information at each site was recovered by sequencing and classifying the distribution of bases synthesized in each barcoded population. Using this method, we successfully decoded 10 out of 11 characters. This proof of concept demonstrates a novel data encoding modality, showing that information can be encoded and decoded into base distributions of DNA in cellular contexts with high accuracy. Unlike previously described cellular encoding systems our approach obviates the need for donor plasmid DNA or pegRNAs^16,17^ and makes the encoding transcription independent and post-translational. We believe the salient parameters of a single effector molecule combined with transcription independent signal capture can unleash a future of new tools for better understanding fast cellular events^18,19^.

## Materials and Methods

### Plasmids and cloning

All PCR reactions were performed using Takara PrimeStar Max DNA Polymerase using primer sequences noted in Table S1 unless otherwise noted. Primers and gBlocks were obtained from Integrated DNA Technologies (IDT). Gibson assemblies were performed using the NEB Gibson Assembly Master Mix according to the manufacturer’s protocol. Unless otherwise noted, plasmid cloning and maintenance were performed in *E. coli* DH5α. Bacterial cultures were maintained in LB Miller Broth (10 g/L casein peptone, 5 g/L yeast extract, 10 g/L NaCl) supplemented with appropriate antibiotics.

Plasmids used in this study are noted in Table S2. hgRNA-D21_pLKO-Hyg (Addgene Plasmid #100562), hgRNA-E21_pLKO-Hyg (Addgene Plasmid #100563), and hgRNA-A21_pLKO-Hyg (Addgene Plasmid # 100559) were ordered from Addgene^6^. pcDNA-Cas9-T2A-TdT (Addgene Plasmid #126424) and pcDNA-Cas9-T2A-STOP-TdT (Addgene Plasmid #126425) were ordered from Addgene^9^. hgRNA-D21-BCLib_pLKO-Hyg was constructed by Gibson assembly of a gBlock encoding the D21 spacer sequence and hgRNA scaffold and a 10 bp degenerate site downstream of the U6 terminator (Fig S1). To isolate individually barcoded variants, hgRNA-D21-BCLib_pLKO-Hyg was transformed into NEB Stable *E. coli* cells and plated on LB agar supplemented with 100 μg/mL ampicillin. Individual colonies were isolated, and the integrity and uniqueness of the barcoded sites was assessed by Sanger sequencing. Barcode sequences isolated and used in this study are compiled in Table S3.

### Cell Culture

All cells were cultured at 37 °C and 5% CO_2_. HEK293T cells and hgRNA cell lines were maintained in DMEM, high glucose, GlutaMAX Supplement, HEPES (Gibco 10564011) supplemented with 10% FBS (Gibco A3160401) and 50 U/mL penicillin and 50 μg/mL streptomycin (Gibco 15070063). Lenti-X 293T (Takara 632180) cells were maintained in DMEM, high glucose, pyruvate (Gibco 11995040) supplemented with 10% FBS (Gibco A3160401).

### Genomic integration of hgRNA sites

Approximately 0.3×106 lenti-X cells per well were seeded into 6-well plates and cultured to 70% confluency. Lentivirus was produced using a second-generation packaging system; each well was transfected with 380 ng psPAX2 (Addgene Plasmid #12260), 140 ng pMD2.G (Addgene Plasmid #12259), and 480 ng of pLKO (Addgene Pladmid #10878) transfer plasmid using 10 μL Lipofectamine 2000 (Thermo Fisher 11668) according to the manufacturer’s protocol. For barcoded cell lines, lentivirus was prepared using aa hgRNA-D21-BCLib_pLKO-Hyg transfer plasmid with a unique barcode sequence for each barcoded variant. For the AED21-hgRNA cell line, three preparations of lentivirus were made using hgRNA-D21_pLKO-Hyg, hgRNA-E21_pLKO-Hyg, and hgRNA-F21_pLKO-Hyg transfer plasmid. After 12 h, the growth medium was exchanged for fresh media, and the cells were cultured an additional 48 h. Lentivirus was harvested by collecting the media from each well and centrifuging for 2 minutes at 500 *x*g. The supernatant was filtered through a 0.45 μm filter and stored on ice at 4°C. Lentivirus was used within 3 days of isolation.

HEK293T cells were passaged in to 6-well plates and cultured to 70% confluency. For barcoded cell lines, each well was transduced with 350 μL of the barcoded lentivirus and 1 μg/mL polybrene. For the AED21-hgRNA cell line, the cells were transduced with 100 μL each of A21, D21, and E21 lentivirus and 1 μg/mL polybrene. After 24 h, the media was refreshed and supplemented with 200 μg/mL hygromycin. Cells were cultured for approximately 2 weeks under hygromycin selection before any subsequent experiments. Media was supplemented with hygromycin for the duration of all experiments. The barcode sequence of barcoded hgRNA sites was validated by deep sequencing.

### hgRNA site extension and nucleotide precursor treatment

Approximately 0.05×10^6^ AED21-hgRNA HEK293T cells were seeded to a poly-D lysine coated 24-well plate and cultured to 70% confluency. Cells were transfected with 500 ng of pcDNA-Cas9-T2A-STOP-TdT (for NHEJ-mediated additions) or pcDNA-Cas9-T2A-TdT (for TdT-mediated additions) using 2 μL Lipofectamine 2000 according to the manufacturer’s protocol. Approximately 4 h after transfection, the media was replaced with fresh media supplemented with nucleotide precursors to the noted final concentration. Working stock solutions were prepared to the following concentrations: 10 mM dGuo (2’-Deoxyguanosine monohydrate, Sigma D7145), 50 mM dThd (2’-Deoxythymidine, Sigma T1895), 100mM dCyd (2’-Deoxycytidine, Sigma D3897), 3mM dAdo (2’-Deoxyadenosine monohydrate, Sigma D7400). Stock solutions were prepared fresh for each experiment in nuclease-free water or DMSO, depending on nucleoside solubility. Cells treated with dAdo were simultaneously treated with dCF (Deoxycoformycin, Sigma SML0508) to a final concentration of 3 μM (add citation). Cells were cultured for 72 h after initial treatment, with media replaced every 24 h. After 72 h, cells were trypsinized, resuspended in 1 mL DMEM, and washed once with PBS. Cells were pelleted, decanted, and stored at -80°C.

### Cellular data encoding

Alphanumeric data was converted into a series of paired nucleotide treatment conditions using the encoding table (Fig 2A). Barcodes were assigned to conditions randomly. Barcoded cell lines were treated with the assigned conditions according to the protocol above, using barcoded hgRNA cell lines and the treatment concentrations in Figure 2B. In parallel, a set of control cells were treated with each of the encoding conditions in triplicate. Cells were trypsinized after 72 h of culturing with the assigned media treatment condition and resuspended in 1mL DMEM. An 850 μL aliquot of each suspension was washed once with PBS and stored at -80°C for subsequent DNA recovery.

### DNA library preparation and sequencing

Sequencing methods were adapted from Kalhor *et al*^6^. Genomic DNA was recovered from frozen cell pellets using the Qiagen DNeasy Blood & Tissue kit according to the manufacturer’s protocol. Recovered genomic DNA was eluted in nuclease free water. The hgRNA locus was amplified using the appropriate primer pair as noted in Table S1 using the Biorad iQ SYBR Green Supermix and approximately 4-8 ng of the genomic DNA. Reactions were amplified by an initial 3 min, 95°C denaturing step followed by cycles of 10 s at 95 °C and 30 s at 60°C. Reactions were monitored in real time and cycling was stopped at mid-exponential amplification, typically 25 cycles. The products of the first reaction were used as a template for a second amplification with NEBNext Dual Indexing Primers. Each sample received a unique index. Reactions were amplified using the same cycling conditions until mid-exponential phase, typically 7 cycles. An equal volume of each sample was pooled into a QC pool. The length distribution and concentration of the library was determined using an Agilent 4200 Tapestation. Pool concentrations were further characterized by Qubit and qPCR methods. Sequencing was performed on an Illumina MiniSeq Mid Output flow cell. Libraries were supplemented with 15-20% phiX control library to increase clustering diversity. After demultiplexing, read counts for each sample were used to re-pool samples for a final sequencing run with evenly balanced indexing across all samples. The balanced pool was sequenced using the methods as before. Library preparation and sequencing were performed at the Rush Universtiy Genomics and Microbiome Core Facility.

### Deep sequencing preprocessing

Illumina sequencing data was preprocessed using a custom python script locally (Windows Surfacebook 2, Intel i7-8650U CPU, 1.90 GHz, 16 GB RAM). The inserted sequence as well as the barcode was extracted from each sequence. The barcode was extracted by searching the reverse (R2) reads for the reverse primer sequence (GCCATACCAATGGGCCCGAATTC) allowing for up to 6 errors (insertions, deletions, and/or substitutions). If this sequence was found, we then searched for the post-barcode sequence (CCTGCAGGAAAAAAA) allowing up to 2 errors (insertions, deletions, and/or substitutions). If both of these sequences were found, the barcode was extracted starting right after the end of the reverse primer sequence and ending at the base preceding the post-barcode sequence. Because these are reverse reads, the barcode was reverse complemented to obtain the final barcode sequence.

The inserted sequence was also extracted from each read. This was done by searching for the D21 hgRNA reverse complement sequence (CTTGGCCGTAGCGTGAC) in the reverse (R2) reads, allowing for no errors. We then searched for TCTAACCCCAC, which is the post-insert sequence directly after the cut site in the R2 reads, allowing for no errors. We then extracted the sequence (if any) inserted between the hgRNA sequence and this post-insert sequence. This insert was reverse complemented to obtain the final insert sequence. This resulted in a set of insert-barcode pairs for each sample.

For the mock pool, all insert-barcode pairs across all control samples were combined into a single pool before All insert-barcode pairs for all encoded samples were combined into a separate pool. This resulted in two mock pools: a mock control pool and a mock encoded pool.

### Cellular data decoding

All decoding was performed locally (Windows Surfacebook 2, Intel i7-8650U CPU, 1.90 GHz, 16 GB RAM). The unpooled samples and mock pooled samples were decoded using the same workflow, except for how the insert sequences were assigned to either (1) a nucleotide treatment condition (for control samples) or (2) each position in the encoded message (for encoded samples). For the unpooled samples, insert sequences were assigned a condition/position based on the associated sample (i.e., based on filename of the fastq file for those sequences). For the mock pooled samples, insert sequences were assigned a condition/position based on the barcode extracted from the same full sequence (Table S4). No insertion, deletion, or substitution errors were allowed when matching sequences to condition/position based on barcode.

Next, the average length and nucleotide compositions were calculated across all sequences for each control condition, *c*, (*X*_*cA*_,*X*_*cC*_, *X*_*cG*_, *X*_*cT*_) or message position, *m* (*X*_*mA*_,*X*_*mC*_,*X*_*mG*_,*X*_*mT*_). When comparing nucleotide compositions for each message position, *m*, against those for each control condition, *c*, we cannot perform most statistical tests as we would violate the principle of normality due to the total sum rule (all elements of the composition add up to 100% so are not independent). Thus, the data is first transformed by using the center log-ratio (clr) transformation which maps this 4-component composition from a 3-dimensional space to a 4-dimensional space, as done in Bhan *et al*^15^ (Equation 1). This is also known as Aitchison space^20^.

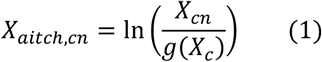

Here, *X*_*cn*_ is composition of nucleotide, *n*, at control condition, *c*. *X*_*aitch,cn*_ is the respective *X*_*cn*_ converted into Aitchison space. *g*(*X*_*c*_) is the geometric mean for condition, *c*, across all four bases in *N* = {*A,C,G,T*} (Equation 2).

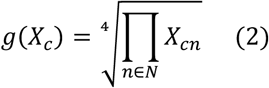

Analogous equations are used for message position, *m*, rather than control condition, *c*:

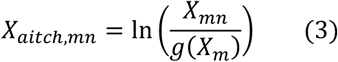

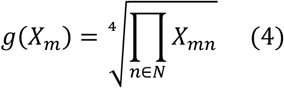

To decode the condition at each position in the encoded samples, we performed statistical tests between the encoded and control samples to calculate the likelihood of each of the six possible conditions, *c* ∈ {1,2,3,4,5,6}. Specifically, we created five probability density functions (PDFs) for each of the six control conditions. Each control condition’s PDFs were generated as normal distributions with the mean and standard deviation of that condition’s three replicates for the respective variables, average length (*L*_*c*_) and the four nucleotide compositions in Aitchison space (*X*_*aitch,cn*_). Next, the likelihood of each encoded sample’s average length and four base compositions were calculated for each of the six conditions, *c*, at each message position, *m*. To choose the most likely condition at each message position, we used Equation 5 to calculate the final likelihood, *P*_*f*_(*c,m*), at each condition, *c*, and message position, *m*, equally weighting the average length PDF value (*P*_*L,c*_(*L*_*m*_)) with the sum of the PDF values for each nucleotide, *n* (*P*_*X,cn*_, *N* = {*A,C,G,T*}).

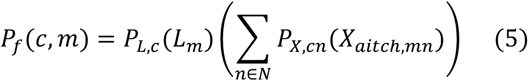

The condition, *c* ∈ {1,2,3,4,5,6}, with the highest *P*_*f*_(*c,m*) was then determined to be the decoded condition for each message position, *m*. Each *P*_*f*_(*c,m*) was divided by the maximum *P*_*f*_(*c,m*) for that position, *m*, to calculate a final normalized value between 0 and 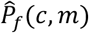 (Equation 6). The encoding table (Table 2A) was then used to decode the entire message based on these decoded condition numbers.

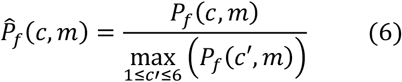

Additionally, reads were subsampled from mock pool samples and Equation 6 was used to decode the message for each subsample. To account for variability in subsampling, multiple replicates were performed by varying the random seed. Specifically, five conditions were tested: (1) all reads (153k for control pool, 244k for message pool) for 1 replicate, (2) 100,000 reads for each pool for 5 replicates, (3) 10,000 reads for each pool for 15 replicates, (4) 1,000 reads for each pool for 30 replicates, and (5) 100 reads for each pool for 50 replicates.

## Results

### TdT catalyzed dNTP additions at hgRNA sites are dependent on the intracellular nucleotide pool

To investigate the nucleotide composition of TdT-catalyzed genomic additions *in vivo*, we used HEK293T cells hosting three previously characterized homing CRISPR-Cas9 sites^6^. Homing guide RNA are designed to contain a protospacer adjacent motif (PAM) between the spacer and the scaffold resulting in Cas9 being targeted to the genomic locus expressing the guide RNA itself. This allows sequential double stranded breaks at this sites and thus longer TdT-catalyzed base additions^9^. Cells were transfected with a plasmid either expressing just functional SpCas9 (the start codon of the TdT gene was replaced with a stop codon in this construct) or both SpCas9 and TdT. After 72 hours of culturing, genomic DNA was extracted from the cells and the hgRNA sites were selectively sequenced by Illumina amplicon sequencing. Additions made at hgRNA sites were analyzed for nucleotide composition and length of the inserts (Fig 1, Figure S2). In the absence of TdT expression, approximately 5% of reads showed nucleotide additions while in the presence of TdT approximately 19% of the reads showed additions (Figure S3). Moreover, both nucleotide composition and lengths of the insertions were significantly altered upon coexpression of TdT with SpCas9, (with frequency of dATP incorporation increasing from 0.18 to 0.39 and insert length increasing from approximately 1.8 nts to 3.2 nts) (Fig 1B, C). Insertions made in the absence of TdT, likely catalyzed by non-homologous end joining (NHEJ) repair, were predominantly adenosines, consistent with previous observations^21^. In contrast, insertions made in the presence of TdT exhibited a distinct composition of primarily C and G, again consistent with previous observations of TdT additions in cellular contexts (Figure S2)^9,22,23^. After confirming that TdT added nucleotides to genomic DNA in a cellular context, we then tested the effect of perturbing intracellular nucleotide pools on insertion composition.

**Figure 1:**
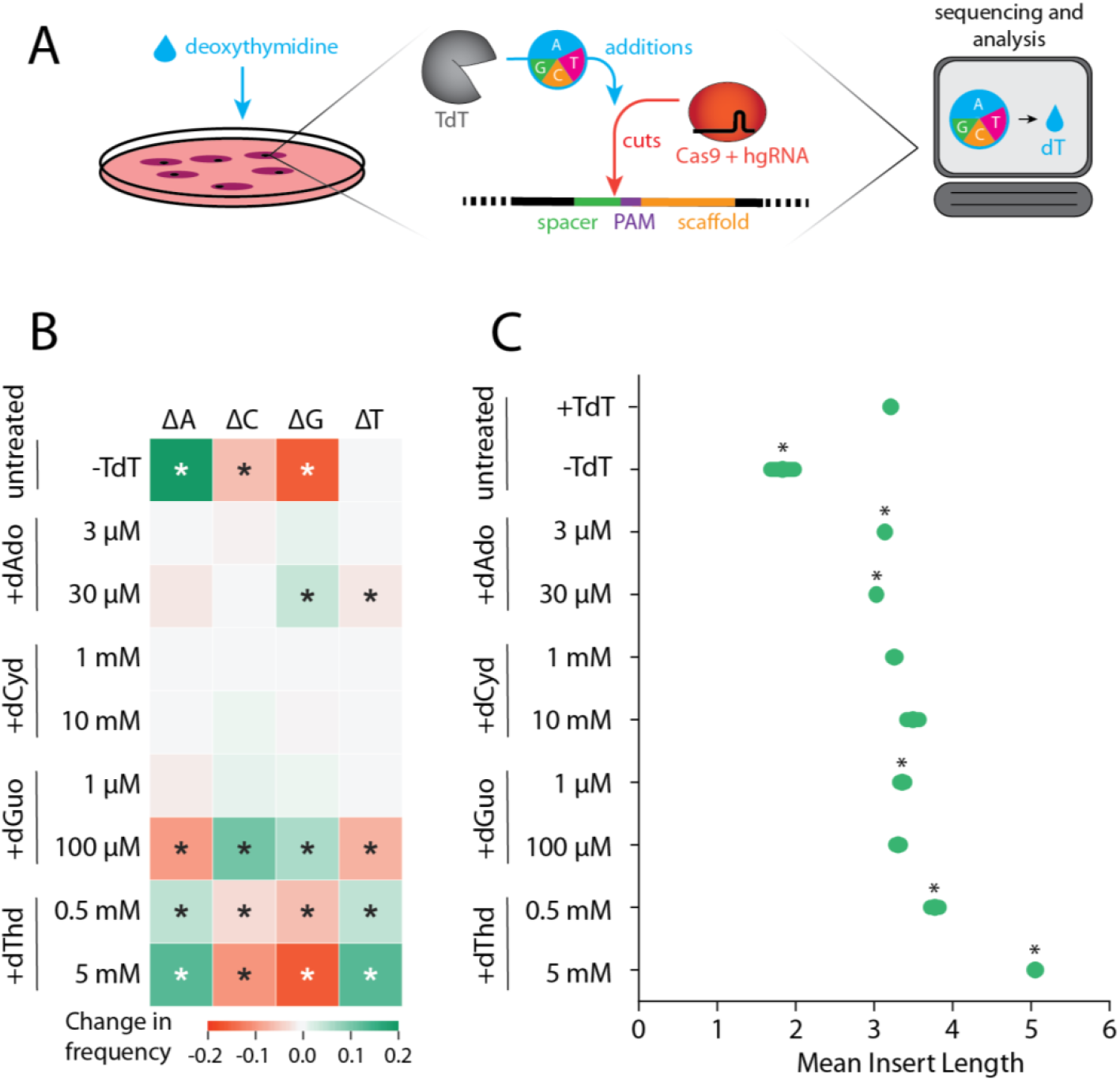
The composition and lengths of inserted nucleotide sequences significantly change in the absence of TdT and upon nucleoside supplementation. (A) Schematic representation of digital data recording experiments. Cells expressing hgRNA, TdT and Cas9 are treated with various nucleosides. Resulting in altered cellular nucleotide pools. This altered condition is encoded into the hgRNA sites by TdT catalyzed DNA strands. Computational analysis of bases added by TdT at cut sites across a population of cells allows *post hoc* inference of the original treatment condition. (B) The composition of inserted sequences changes across conditions (rows), including a large change in the absence of TdT (top row) as well as different nucleoside environments, especially when supplementing dGuo and dThd (last 4 rows). The four columns represent the changes in the four base (A, C, G, and T) frequencies. The color of cells represents the change in frequency of each nucleotide (column) in inserted sequences across conditions (rows) compared to the +TdT condition with no nucleoside treatment (e.g., the inserted dATP frequency increased from 0.18 in the +TdT condition to 0.37 in the -TdT condition, resulting in an increase of 0.19, which is the plotted value for -TdT and ΔA). See Fig S2 for absolute A, C, G, and T frequency values for each condition. Positive changes are shown in green and negative changes are shown in red. Asterisk symbols are displayed for statistically significant changes (compositions were first transformed into Aitchison space before performing two-sample independent t-test with Bonferroni correction: α < 0.05/36 = 0.00139). (C) The mean length of inserted sequences in each sample decreases in the absence of TdT (first and second row) and varies across nucleoside treatments (all other rows). Green bars are one standard deviation on either side. Asterisk symbols are displayed for statistically significant changes in mean length from the +TdT condition (first row) (two-sample independent t-test with Bonferroni correction: α < 0.05/9 = 0.0056). Zero-length inserts were not included in mean length calculations (see Fig S3 for insertion rates).

Gangi-Peterson *et al*. previously reported that changes in the base composition of TdT-mediated additions at V(D)J junctions depend on intracellular dATP pools^22^. Thus, we hypothesized that altered nucleotide pools would similarly bias the composition of DNA synthesized by TdT at hgRNA sites. Cells hosting hgRNA sites were transfected with a plasmid coexpressing SpCas9 and TdT. Cellular nucleotide pools were altered by daily treatment with deoxyribonucleosides (dNs: deoxyadenosine (dAdo), deoxycytidine (dCyd), deoxythymidine (dThd), or deoxyguanosine (dGuo)) for the 72-hour culture period. Cells were treated with the adenosine deaminase inhibitor, dCF, for 20 minutes before dAdo addition to prevent conversion of dAdo to deoxyinosine^22,24^.Upon extracellular dosing with dNs, we observed significant changes in the composition of additions at the hgRNA sites in a dose dependent manner, with the exception of dCyd which resulted in no significant difference at the concentrations tested (Fig 1B). One of the potential reasons for poor response to dCyd perturbation could be that dCTP has been shows to be the least favored TdT substrate *in vitro*, especially if the 3’ OH base is a dC^25^. We observed a significant increase in G incorporation and a decrease in T incorporation frequency upon treatment with 30 μM dAdo, and a significant increase in G and C incorporation frequencies upon treatment with 100 μM dGuo (Fig 1B). We also observed a significant increase in A and T incorporation frequencies when the cells were dosed with 0.5 mM dThd, with an even larger shift occurring when treated with 5 mM dThd. After observing these significant shifts in nucleotide composition, we sought to employ these cellular states to encode digital information.

### Post-translational digital data encoding into genomes of mammalian cell population with high accuracy

Having established which cellular nucleoside perturbation states could be encoded at the hgRNA sites we identified conditions that would maximize the differences in the distribution of bases added (Fig 2B). We tested 0.025, 0.05 and 0.1 mM dGuo, and ascertained that 4 nucleoside treatment conditions (0.025, 0.1 mM dGuo, 0.5 and 5 mM dThd) could be mutually differentiated from each other and from an untreated condition. Additionally, we determined that we could differentiate between conditions with and without TdT expression largely due to a significant difference in insertion length (Fig 1C). Collectively, we found 6 differentiable conditions (Fig 2B, S4, S5), and devised an alphanumeric encoding scheme (Fig. 2A, B) allowing a maximum of approximately 2.6 bits to be encoded in a single population of hgRNA expressing cells.

**Figure 2:**
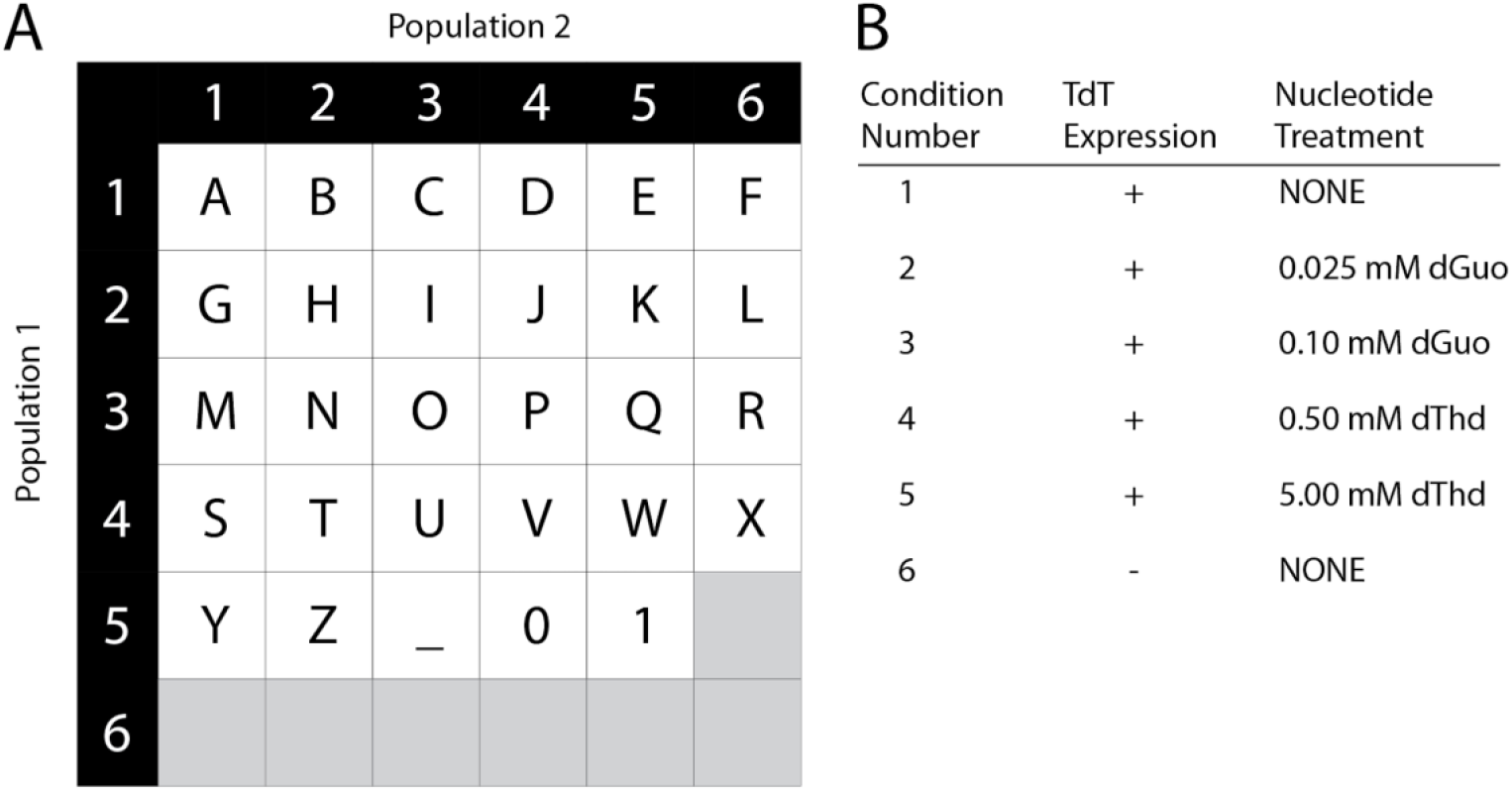
Data encoding scheme and barcoded locus design. (A) Encoding table for representing data. Each character is encoded via two cell populations, each population encoded as one of six conditions (Fig 2B). (B) Six distinguishable conditions are used for data encoding with varied TdT expression and nucleoside treatment.

To expand the encoding capacity further, we reasoned that we could barcode cellular populations so that a combination of two barcoded populations act as a single memory address. To accomplish this, we created DNA cassettes that encoded both hgRNA expression and specified barcodes, packaged them into lentiviral particles and generated 24 populations of HEK293T with unique barcodes (Fig. S6). In this case, two barcoded populations when treated with two nucleoside perturbation states would result in the encoding of a single alphanumeric character at a particular position in our digital information.

As a proof of concept for composition-based data encoding, we assigned a single character from the message “HELLO WORLD” to ordered pairs of memory addresses. Cell populations were transfected and treated according to the assigned conditions (Fig. 3A). The cells were harvested, and barcoded hgRNA sites were sequenced as before. Individual sequencing reads were assigned to a memory address using indexing barcodes or locus barcode sequences. For each memory address, the length and composition of additions made at the corresponding hgRNA sites was calculated, and a custom likelihood function was used to estimate the most probable treatment condition (Equation 5). 21/22 addresses were correctly decoded, yielding the message “HELLO WORLV” (Fig. 3A). While one of the values was incorrectly decoded, the correct value was the second most likely (Fig. 3A).

**Figure 3:**
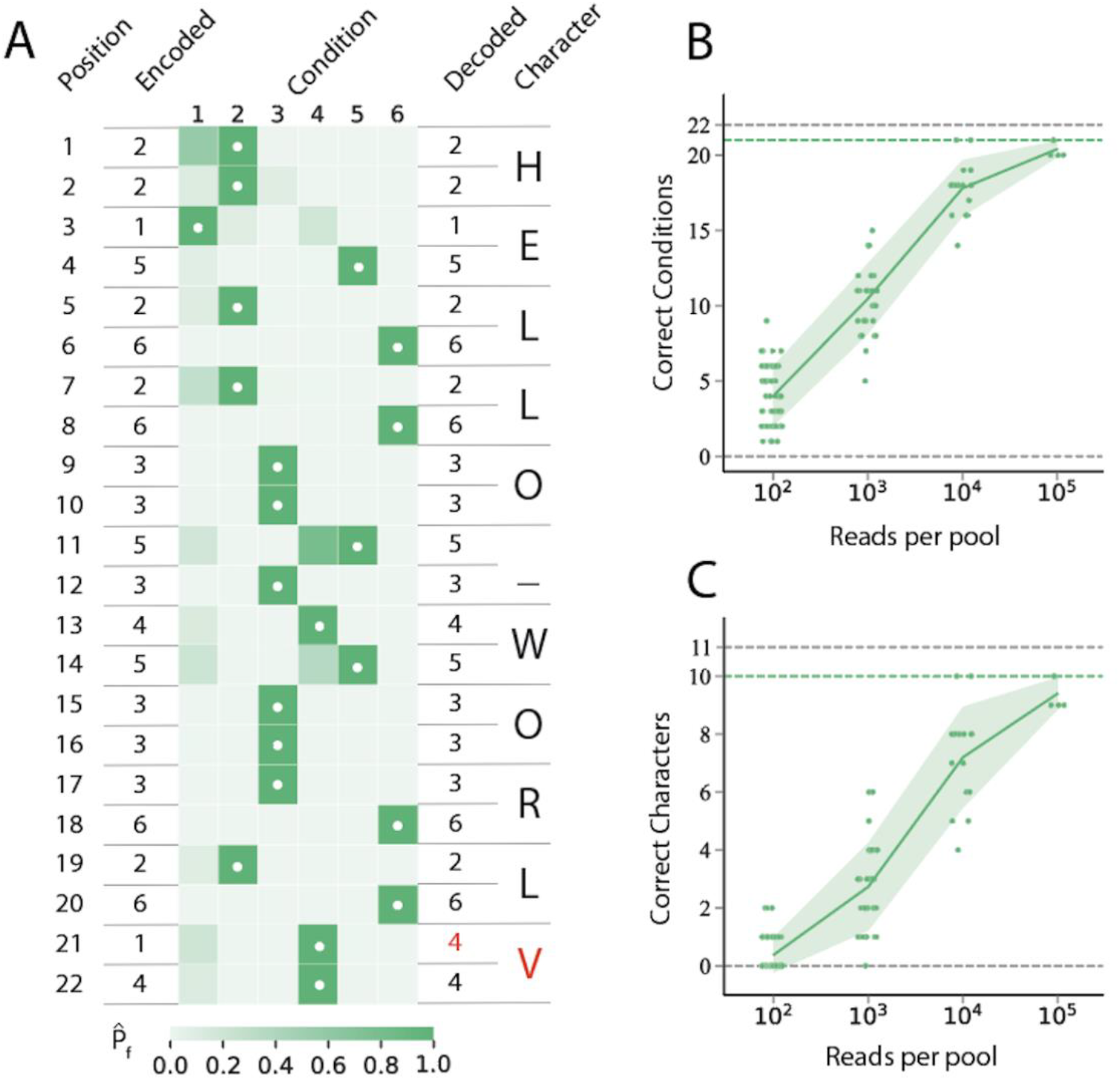
The encoded message, “HELLO WORLD”, was decoded with one error as “HELLO WORLV”. Decoding was performed again after subsampling insert sequences *in silico* to assess performance with less data. (A) “HELLO WORLD” was encoded as a string of 22 numbers (two per character, including a space). The length and nucleotide composition of the inserts of each encoded sample were compared to controls for each condition to calculate 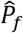 the normalized likelihood of that condition (Equations 5 and 6). 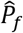 values are shown in the heatmap. White dots indicate the decoded condition (highest 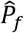). The table in Figure 2A was used to decode each ordered pair of decoded values into characters shown at the right. One error was found at position 21 (red value). (B, C) Insert sequence data was repeatedly subsampled *in silico* to estimate the impact that the quantity of sequence data had on decoding performance. Decoding performance was measured as the number of correctly decoded conditions (B) or correctly decoded characters (C). The control pool and message pool were each subsampled to have the number of reads shown on the x-axis. Each green dot shows the performance of a different (random) subsample. Some horizontal jitter is applied to these values to better visualize the distribution. The solid green lines show the mean values with green shading extending one standard deviation on either side. Dotted gray lines denote the required value for 100% decoding accuracy and dotted green lines show the value obtained with all data (roughly 153k reads for the message pool and 244k for the control pool).

Next, we investigated the effect of read counts on decoding accuracy. Specifically, sequencing reads for all samples were pooled and subsampled to generate smaller datasets. Multiple replicates were performed to account for variability in subsampling. Again, the message was decoded from the subsampled pools using hgRNA locus barcodes and the number of correctly decoded addresses/characters was calculated (Fig. 3B, C). We found that having greater than 10^5^ reads for both the control and message pools resulted in similar performance as using all reads. Overall, this shows that information can be encoded in the genome as nucleotide compositions generated via TdT.

## Discussion

Recording cellular signals that alter at minutes’ time scale with high temporal resolution can only be achieved by employing a post-translational enzymatic recording system. As a first step towards achieving this we theorized we needed to prototype a recording system where the enzymatic effector molecule is independent of signal-induced expression. In various mammalian cells average transcription time is 10 minutes/gene(for a 10kb gene, average size of mammalian genes) and the average translation time is 1 minute/300aa protein^26^. On the other hand, an expressed protein can last in the cell for about 24 hours while a metabolite of interest lasts only about 1 minute, depending on the identity of the metabolite^26^. The average turnover number for TdT is between 0.0925-1.2 s^-1^, and TdT has been shown to be able to record various biologically relevant molecules^27^. Thus, deploying TdT as the effector molecule a post-translational signal recording system possible.

To better understand the characteristics of such a system, we encoded predetermined digital information into the genomes of mammalian cells. We were able to do so with high accuracy (91%), using just 6 treatment conditions (“signals”).

While we used only 4 nucleoside treatment conditions for encoding, using larger and/or intermediate concentrations of each nucleoside could increase the data encodable at each barcoded locus as well as decoding accuracy. Additionally, TdT can incorporate unnatural nucleotides^28–31^ which would significantly expand the set of accessible nucleotide compositions. Altering intracellular nucleotide pools for information storage has the advantage of being highly multiplexable: except for dCyd, each nucleoside yields a distinct DNA composition in a dose-dependent manner with no modifications of the encoding apparatus (TdT+Cas9+hgRNA locus) itself.

This system also shows potential in physiological applications. While modifications to intracellular nucleotide pools using nucleoside treatment was a convenient proof of concept, imbalanced nucleotide pools may also arise physiologically in neurological disorders and immunodeficiencies that result from errors in nucleotide metabolism^32,33^. Applied to model organisms, the method described here may be useful as a means to characterize nucleotide imbalances *in situ* among heterogenous cell populations.

By coupling the catalytic characteristics of TdT(s) to other signals of interest, information about cellular states other than nucleotide imbalance could also be encoded in DNA composition. Previously, we have shown that the composition of DNA synthesized by TdT could be modulated by using *in vitro* reactions with two distinct TdT variants with different nucleotide selectivity: one unmodified, and the other engineered to deactivate in the presence of calcium. A similar approach could be used to engineer a two-polymerase system responsive to alternative inputs by fusing sensing domains to TdT that respond to inputs such as light^34^, small molecules^35^, or temperature^36^.

This single enzyme dependent DNA synthesis-based *in vivo* encoding system circumvents the need for transcription and translation, making its response to biosignals inherently faster than previously described systems^3,7–10,12,14,16,17^. We hope this proof of concept will pave the way for development of similar post-translational systems making high resolution recording of cellular biosignals and metabolic states possible.

## Acknowledgements

The authors would like to acknowledge Dr. Neha Kamat and Dr. Curt Horvath for generously sharing cell culture equipment and space. Haley Edelstein provided helpful discussion regarding lentivirus production and transduction. Dr. Jean-Patrick Parisien provided helpful discussion regarding cell culture. Real time PCR was supported by the Northwestern University Keck Biophysics Facility, a shared resource of the Robert H. Lurie Comprehensive Cancer Center of Northwestern University supported in part by the NCI Cancer Center Support Grant #P30 CA060553. Illumina sequencing was performed with the help of the Next Generation Sequencing Core Facility at University of Illinois at Chicago and the Rush Genomics and Microbiome Core Facility. Sanger sequencing was supported by the Northwestern University NUSeq Core Facility. This work was funded by the National Institutes of Health grants R01MH103910 and UF1NS107697 and National Institutes of Health Training Grant (T32GM008449) through Northwestern University’s Biotechnology Training Program (to JS and AC).

