## Supplementary Information for "Post-translational digital data encoding into the genomes of mammalian cell populations"

**Table S1:** primer and gBlock sequences

| Primer | Sequence | Notes |
|---|---|---|
| seq F | ATGGACTATCATATGCTTACCGT | hgRNA site amplification primer F |
| seq R | TTCAAGTTGATAACGGACTAGC | hgRNA site amplification primer R, non barcoded sites |
| seq R bc | GCCATACCAATGGGCCCGAA | hgRNA site amplification primer R, barcoded sites |
| D21 bc gBlock F | gactatcatatgcttaccgtaacttgaaagtatttcgatttctt | Gibson assembly primer for D21 bc gBlock, F |
| D21 bc gBlock R | tgtctcgaggtcgagaattcCGTATAATCATGTGCAGTGT | Gibson assembly primer for D21 bc gBlock, R |
| pLKO-Hyg F | ACACTGCACATGATTATACGgaattctcgacctcgagaca | Gibson assembly primer for pLKO-Hyg backbone, F |
| pLKO-Hyg R | aatcgaaatactttcaagttacggtaagcatatgatagtcca | Gibson assembly primer for pLKO-Hyg backbone, R |
| D21 bc gBlock | aacttgaaagtatttcgatttcttggctttatatatcttgtggaaaggac<br>gaaacaccgGTCACGCTACGGCCAAGGTGgggttagagctagaaatagca<br>agttaacctaaggctagtccgttatcaacttgaaaaagtggcaccgagtc<br>ggtgctttttttCCTGCAGGNNNNNNNNNNGAATTCGGGCCCATTGGTAT<br>GGCGTGCGAAGGAGTGGCTGGACTAATCCGCCAAGCGTATGCGAAGGGTA<br>GAATCATTAGAGTGCTGTACATTTATAAGTTGACTACACTGCACATGATT<br>ATACG | |

**Table S2:** plasmids

| Plasmid Name | Cloning AbR | Source | Notes |
|---|---|---|---|
| hgRNA-A21_pLKO-Hyg | Amp | Kalhor, R., Mali, P., & Church, G. M. (2017). Rapidly evolving homing CRISPR barcodes. Nature Methods, 14(2), 195–200. https://doi.org/10.1038/nmeth.4108 | |
| hgRNA-D21_pLKO-Hyg | Amp | | |
| hgRNA-E21_pLKO-Hyg | Amp | | |
| pcDNA-Cas9-T2A-TdT | Amp | Loveless, T. B., Grotts, J. H., Schechter, M. W., Forouzmand, E., Carlson, C. K., Agahi, B. S., Liang, G., Ficht, M., Liu, B., Xie, X., & Liu, C. C. (2021). Lineage tracing and analog recording in mammalian cells by single-site DNA writing. Nature Chemical Biology, 17(6), 739–747. https://doi.org/10.1038/s41589-021-00769-8 | |
| pcDNA-Cas9-T2A-STOP-TdT | Amp | | |
| hgRNA-D21-BC#_pLKO-Hyg | Amp | This study | 24 variants, where # = barcode ID (Table S3) |

**Table S3:** barcode sequences

| ID | Barcode Sequence |
|---|---|
| 1 | AATTTTGCGG |
| 2 | TGACTTTTAA |
| 4 | TAACAGTATG |
| 6* | TTTTTGTGAA, GTTATACTGT, CAACTCGGTC |
| 9 | CAATTGTCAT |
| 10 | GCCGCTCAGT |
| 15 | TATAGCCACC |
| 16 | TGCACGTCAT |
| 17 | GCGATCCCGG |
| 18 | AACCTAAGTT |
| 19 | ACTGCTCCCT |
| 22 | TTTTACAGAT |
| 23 | ACATAAATTA |
| 24 | CCTTCGACTC |
| 29 | TCCGGACTCA |
| 31 | GGTATAAAAA |
| 34 | CGTATTCTCT |
| 35 | GTCCCAAAAA |
| 37 | AGTTTTTCAA |
| 38 | GGTTACACTT |
| 41 | TCTTAGCATT |
| 42 | GTCCTGCTAA |
| 44 | CAATAACCGA |
| 45 | TTCAATTCGA |

*Three unique barcodes were recovered from cells harboring barcode 6

**Table S4:** barcode mapping

| Treatment controls barcode mapping | | | Message encoding barcode mapping | |
|---|---|---|---|---|
| Nucleoside Treatment | TdT +/- | Barcodes | Message Position | Barcodes |
| 0 | + | AATTTTGCGG | 1 | AATTTTGCGG |
| 0 | + | TGACTTTTAA | 2 | TGACTTTTAA |
| 0 | + | TAACAGTATG | 3 | TAACAGTATG |
| g | + | AGTTTTTCAA | 4 | TTTTTGTGAA,GTTATACTGT,CAACTCGGTC |
| g | + | GGTTACACTT | 5 | CAATTGTCAT |
| g | + | TCTTAGCATT | 6 | GCCGCTCAGT |
| gg | + | TTTTTGTGAA,GTTATACTGT,CAACTCGGTC | 22 | TATAGCCACC |
| gg | + | CAATTGTCAT | 7 | TGCACGTCAT |
| gg | + | GCCGCTCAGT | 8 | GCGATCCCGG |
| t | + | TATAGCCACC | 9 | AACCTAAGTT |
| t | + | TGCACGTCAT | 10 | ACTGCTCCCT |
| t | + | GCGATCCCGG | 11 | TTTTACAGAT |
| tt | + | ACATAAATTA | 12 | ACATAAATTA |
| tt | + | CCTTCGACTC | 23 | CCTTCGACTC |
| tt | + | TCCGGACTCA | 13 | TCCGGACTCA |
| 0 | - | AACCTAAGTT | 14 | GGTATAAAAA |
| 0 | - | ACTGCTCCCT | 24 | CGTATTCTCT |
| 0 | - | TTTTACAGAT | 15 | GTCCCAAAAA |
| gg | - | GTCCTGCTAA | 16 | AGTTTTTCAA |
| gg | - | CAATAACCGA | 17 | GGTTACACTT |
| gg | - | TTCAATTCGA | 18 | TCTTAGCATT |
| tt | - | GGTATAAAAA | 19 | GTCCTGCTAA |
| tt | - | CGTATTCTCT | 20 | CAATAACCGA |
| tt | - | GTCCCAAAAA | 21 | TTCAATTCGA |

```
gagggcctatttcccatgattccttcatatttgcatatacgatacaaggctgttagagagataa
ttggaattaatttgactgtaaacacaaagatattagtacaaaatacgtgacgtagaaagtaata
atttcttgggtagtttgcagttttaaaattatgttttaaaatggactatcatatgcttaccgta
acttgaaagtatttcgatttcttggctttatatatcttgtggaaaggacgaaacaccgGTCACG
CTACGGCCAAGGTGgggttagagctagaaatagcaagttaacctaaggctagtccgttatcaac
ttgaaaaagtggcaccgagtcggtgctttttttCCTGCAGGNNNNNNNNNNGAATTCGGGCCCA
TTGGTATGGC
```

U6 promoter
Spacer
Scaffold
U6 terminator
Locus barcode
seq primer binding sites

**Figure S1:** barcoded D21 hgRNA site sequence. See Table S3 for locus barcode sequences.

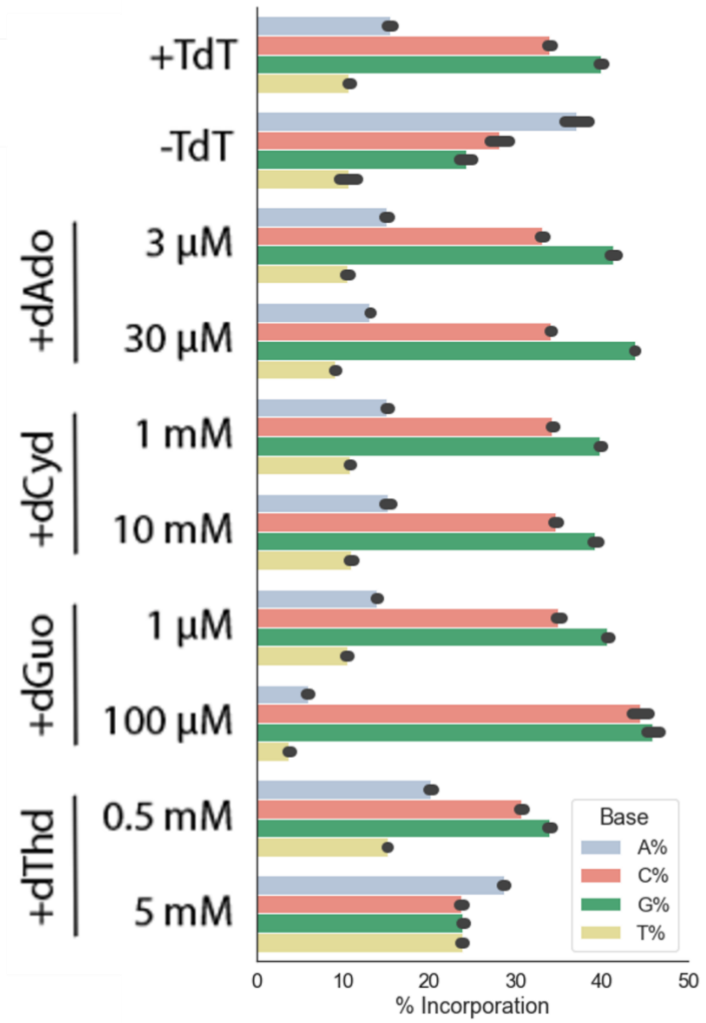


**Figure S2:** Percent incorporation of each base across all reads with additions (at least 1 base incorporated) for each condition. Bar height shows mean value and error bars are one standard deviation above and below the mean value.

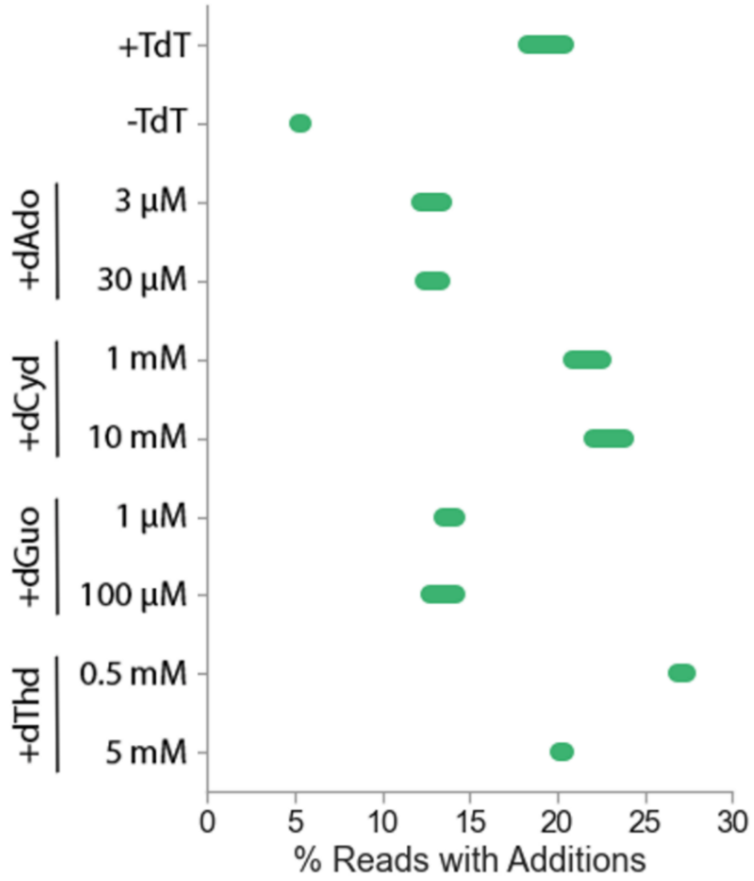


**Figure S3:** The percentage of reads containing hgRNA sites with additions (at least one base incorporated) increases in the presence of TdT (top row vs second row) and varies based on nucleoside treatment (all other rows). Width of each bar is 2 standard deviations (1 standard deviation on each side from mean in middle).

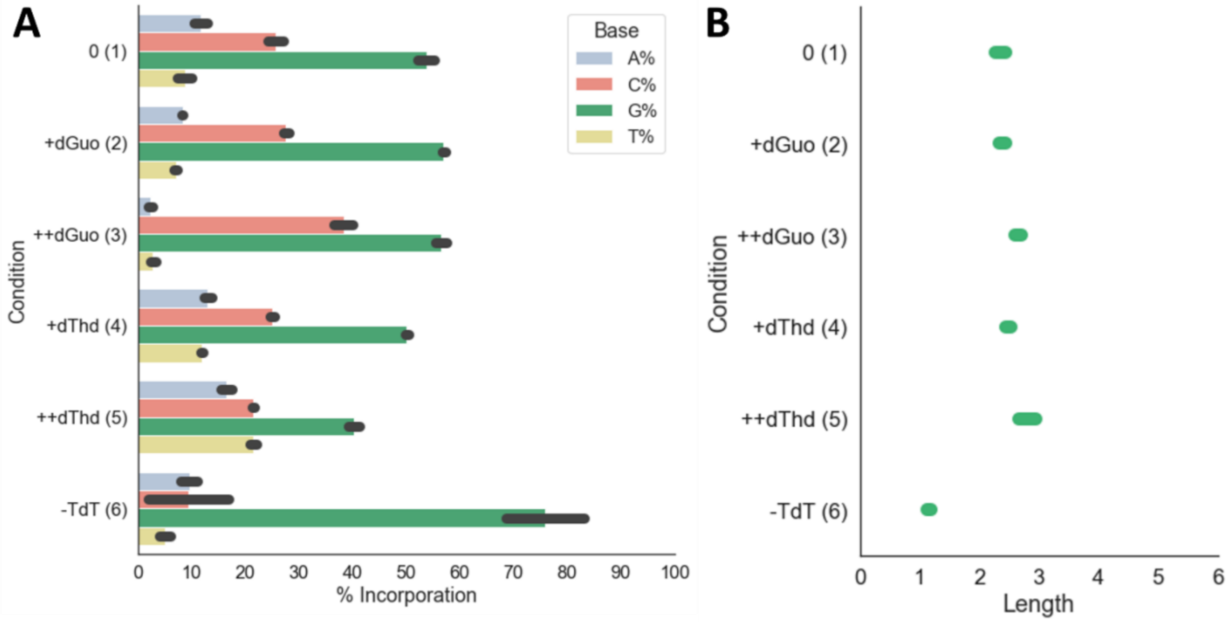


**Figure S4:** (A) Percent incorporation of each base across all reads with additions (at least 1 base incorporated) for each condition in the message encoding experiment. Bar height shows mean value and error bars are one standard deviation above and below the mean value. Condition numbers (1-6) are shown in parentheses (Fig 2B). (B) Average length of additions for each condition.

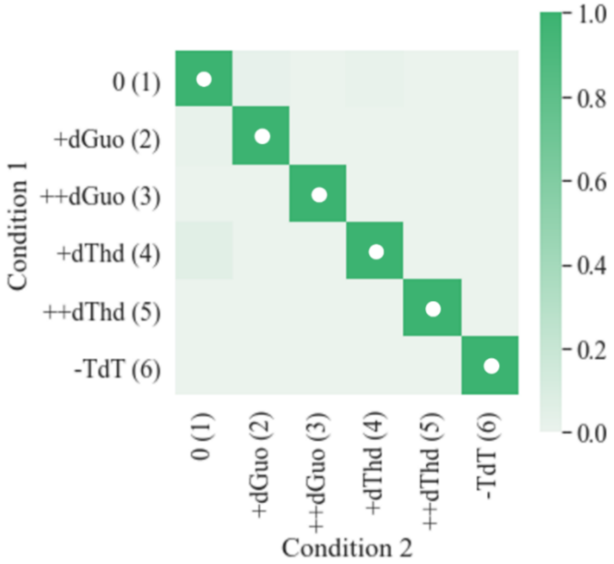


**Figure S5:** Controls are easily distinguished from each other on the basis of length and nucleotide composition. Overlap is seen only for conditions 1 and 4 and is minimal. Color shows normalized likelihood (Equation 6) that the values for a sample from Condition 2 are derived from the Condition 1 length and nucleotide composition PDFs (Equation 5). Condition numbers (1-6) are shown in parentheses (Fig 2B).

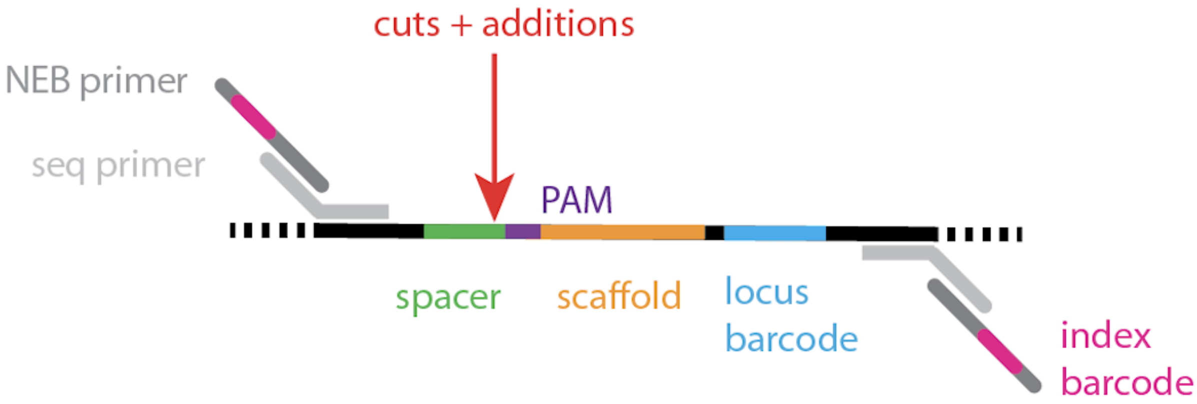


**Figure S6:** Schematic of barcoded hgRNA barcoded loci and sequencing primers. hgRNAs target Cas9 to the spacer and PAM sequence, allowing TdT to make additions. The downstream locus barcode remains constant and enables demultiplexing from pooled data. Memory locus sequences are recovered from genomic DNA by amplification with seq primers and prepared for Illumina sequencing with barcoded NEB primers.
